# Insulin-like growth factor 1 receptor signaling in tenocytes is required for adult tendon growth

**DOI:** 10.1101/670026

**Authors:** Nathaniel P Disser, Kristoffer B Sugg, Jeffrey R Talarek, Dylan C Sarver, Brennan J Rourke, Christopher L Mendias

**Author notes:** These authors contributed equally to this work (Nathaniel P Disser, Kristoffer B Sugg). To whom correspondence should be addressed:* Christopher L Mendias, PhD, Hospital for Special Surgery, 535 E 70th St, New York, NY 10021, USA, office.

## Abstract

Tenocytes serve to synthesize and maintain collagen fibrils and other matrix proteins in tendon. The underlying biological mechanisms of postnatal tendon growth and repair are not well understood. Insulin-like growth factor 1 (IGF1) plays an important role in the growth and remodeling of numerous tissues, but less is known about IGF1 in tendon. We hypothesized that IGF1 signaling is required for proper tendon growth in response to mechanical loading through regulation of collagen synthesis and cell proliferation. We conditionally deleted the IGF1 receptor (*IGF1R*) in scleraxis (*Scx*) expressing tenocytes, and compared to control *Scx:IGF1R*^+^ mice, *Scx:IGF1R*^Δ^ mice demonstrated reduced tenocyte proliferation and smaller tendons in response to mechanical loading. Additionally, we identified that both the PI3K/Akt and ERK pathways are activated downstream of IGF1 and interact in a coordinated manner to regulate cell proliferation and protein synthesis. These studies indicate that IGF1 signaling is required for proper postnatal tendon growth.

## Introduction

Tendon is a dense connective tissue that serves to transmit force from muscle to bone during mechanical loading. The tendon extracellular matrix (ECM) is composed mostly of type I collagen, as well as type III collagen, elastin, and proteoglycans (Gumucio et al., 2015). Tendon fibroblasts, or tenocytes, are the main cell type in tendon and are responsible for the synthesis, organization, and maintenance of the ECM (Wang, 2006). In response to high stress repetitive mechanical loading, such as that which occurs during resistance exercise, tendons adapt by undergoing hypertrophy (Couppé et al., 2008). This results in an increase in tendon cross-sectional area (CSA) through an induction of cell proliferation and collagen production (Geremia et al., 2018; Gumucio et al., 2014; Svensson et al., 2016). Repetitive loading can also induce a chronic inflammatory condition referred to as tendinopathy (Mead et al., 2018; Millar et al., 2017). Despite the high prevalence of tendon injury and overall importance of tendon in maintaining musculoskeletal health, the underlying biological mechanisms that regulate postnatal growth are not well understood. Further, as tendinopathy is thought to arise due to improper responses to mechanical stimuli, gaining a greater understanding of the basic biological mechanisms of tendon growth could help develop new therapies for the treatment of tendon disorders.

Insulin-like growth factor-1 (IGF1), which can be induced by growth hormone (GH) and other mechanical signals, is an integral component of the growth and development of several different tissues (Heinemeier et al., 2012). IGF1 is typically bound to an IGFBP carrier protein, and upon proteolytic degradation of the IGFBP, IGF1 is liberated and can interact with its receptor, IGF1R which is a member of the receptor tyrosine kinase (RTK) family of transmembrane receptors (Lelbach et al., 2005). IGF1 is a potent activator of skeletal muscle cell proliferation and protein synthesis, and the deletion of *IGF1* in muscle fibers results in fiber atrophy and disrupted metabolism (Heinemeier et al., 2012; Vassilakos et al., 2019). In bone, overexpression of IGF1 results in increased bone mineral density, and the inactivation of *IGF1R* in osteoblasts impairs matrix mineralization (Yakar et al., 2010). However, the role of IGF1 in tendon growth has not been fully examined. Previous studies in tendon have revealed an increase in tendon collagen synthesis following IGF1 treatment in engineered tendon tissue and healthy human tendon, as well as in horses with tendinopathies (Dahlgren et al., 2002; Herchenhan et al., 2015; Nielsen et al., 2014b). Given the role that IGF1 plays in the growth of other tissues, and observations of increased IGF1 expression correlated with enhanced ECM production in tendons, we sought to determine the mechanisms behind IGF1-mediated tendon growth. We hypothesized that IGF1 signaling is required for proper tendon growth in response to mechanical loading through a coordinated induction of collagen synthesis and cell proliferation. To test this hypothesis, we induced tendon growth in adult mice via mechanical overload of the hindlimb, and deleted *IGF1R* in scleraxis expressing tenocytes. Additionally, we performed a series of *in vitro* experiments where we treated tenocytes with IGF1 for various durations to examine the molecular mechanisms induced by IGF1 during postnatal tendon growth.

## Results

We generated mice that enabled us to perform a targeted deletion of *IGF1R* in tenocytes by crossing *IGF1R*^flox/flox^ mice in which exon 3 of *IGF1R* is flanked by loxP sites (Dietrich et al., 2000), to *Scx*^CreERT2/CreERT2^ mice in which an *IRES-CreERT2* sequence is present between the stop codon and 3’ UTR in exon 2 of scleraxis (Howell et al., 2017). After performing initial crosses between *Scx*^CreERT2/CreERT2^ and *IGF1R*^flox/flox^ mice, mice were backcrossed until a homozygous state was reached. *Scx*^CreERT2/CreERT2^:*IGF1R*^flox/flox^ mice allowed us to inactivate *IGF1R* in scleraxis expressing cells upon treatment with tamoxifen (referred to as *Scx:IGF1R*^Δ^ mice), while *Scx*^+/+^:*IGF1R*^flox/flox^ mice would maintain the expression of *IGF1R* in scleraxis expressing cells after tamoxifen treatment (referred to as *Scx:IGF1R*^+^ mice). An overview of the alleles used in this study is provided in Figure 1A. Wild type C57BL/6J mice were used for cell culture experiments, while *Scx:IGF1R*^+^ and *Scx:IGF1R*^Δ^ mice were used in whole animal studies.

**Figure 1.**
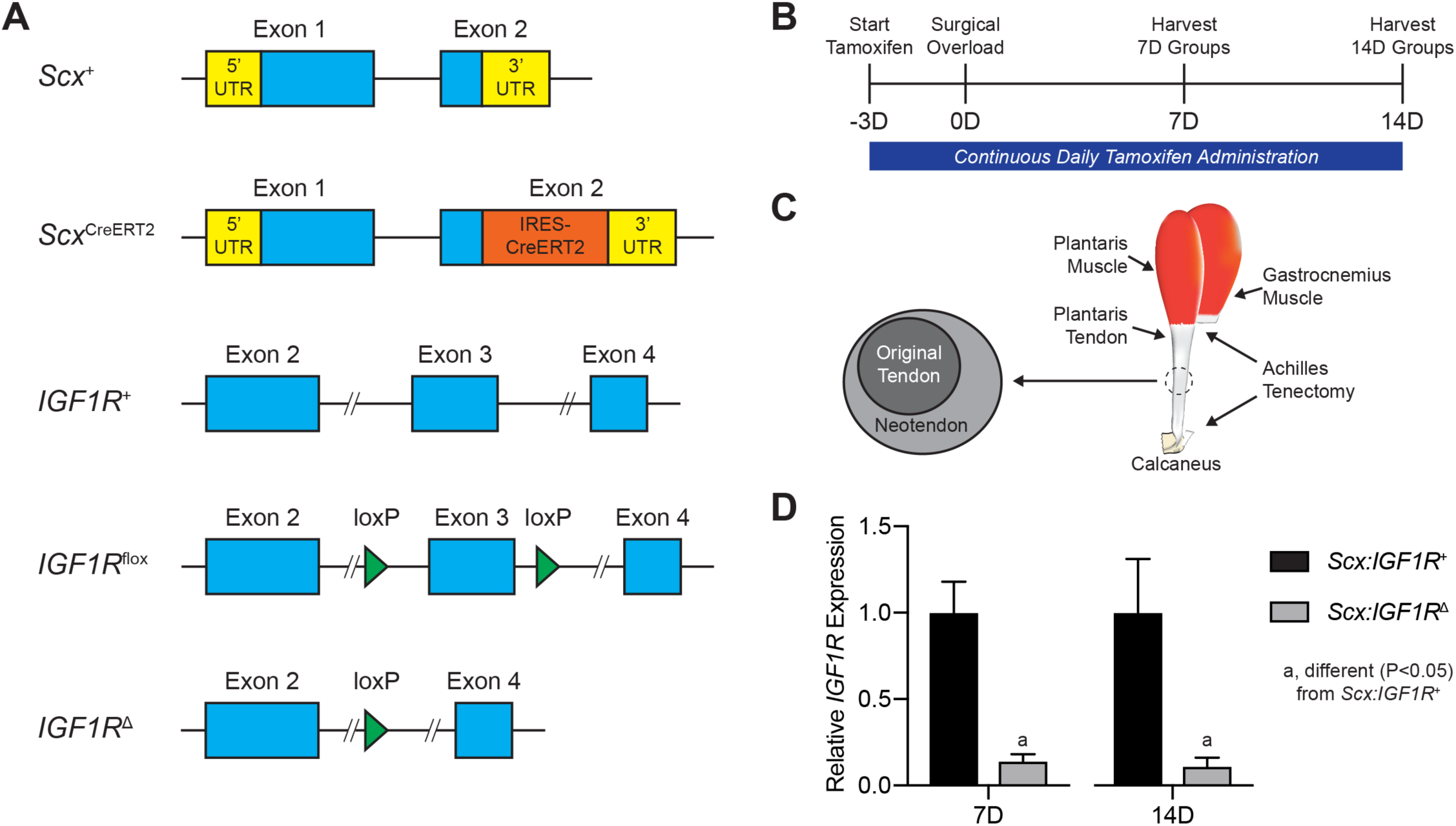
Experimental overview. (A) Overview of the alleles used in this study, including the scleraxis wild type (*Scx*^+^), scleraxis inducible Cre (*Scx*^CreERT2^), IGF1R wild type (*IGF1R*^+^), IGF1R floxed (*IGF1R*^flox^), and IGF1R-null (*IGF1R*^Δ^) alleles. (B) Timeline of tamoxifen treatment, surgical overload, and tissue harvest. Tamoxifen was delivered on a daily basis beginning three days prior to surgery, and continued through tissue harvest. (C) Overview of surgical overload procedure, in which the Achilles tendon is removed from the animal, resulting in compensatory hypertrophy of the synergist plantaris muscle and tendon. A neotendon area of new tendon matrix forms around the original tendon. (D) Relative expression of *IGF1R* at 7 days and 14 days, with the *Scx:IGF1R*^Δ^ group normalized to the *Scx:IGF1R*^+^ group at each time point. Values are mean±CV. Differences for each time point tested with a t-test: a, significantly different (P<0.05) from *Scx:IGF1R*^+^ group. N=4 animals per group.

To study the role of IGF1 signaling in adult tendon hypertrophy, we treated mice with tamoxifen, and then induced a mechanical overload in plantaris tendons of *Scx:IGF1R*^+^ and *Scx:IGF1R*^Δ^ mice, and analyzed tendons at either 7 or 14 days after creation of the growth stimulus (Figure 1A-C). Tamoxifen treatment resulted in an over 90% reduction in *IGF1R* expression in *Scx:IGF1R*^Δ^ mice compared to *Scx:IGF1R*^+^ mice at both 7 and 14 days (Figure 1D). We then analyzed histological changes in plantaris tendons. As previously observed in other models of synergist ablation-mediated mechanical overload, a neotendon tissue formed around the original tendon that matured and filled in with collagen over time, however growth was blunted in the *Scx:IGF1R*^Δ^ mice (Figure 2A-D). The total tendon CSA was not different between groups at 7 days, but by 14 days the total CSA of *Scx:IGF1R*^+^ mice was twice as large as *Scx:IGF1R*^Δ^ mice (Figure 2D). This occurred due to a greater expansion of the neotendon of *Scx:IGF1R*^+^ mice over time, while *Scx:IGF1R*^Δ^ mice displayed no change between 7 and 14 days (Figure 2C). No differences in cell density were observed in the original tendon, neotendon, or total tendon across time or genotype (Figure 2E-G), however the percentage of proliferating cells was two-fold greater in the neotendon of *Scx:IGF1R*^+^ mice compared to *Scx:IGF1R*^Δ^ mice (Figure 2H-J).

**Figure 2.**
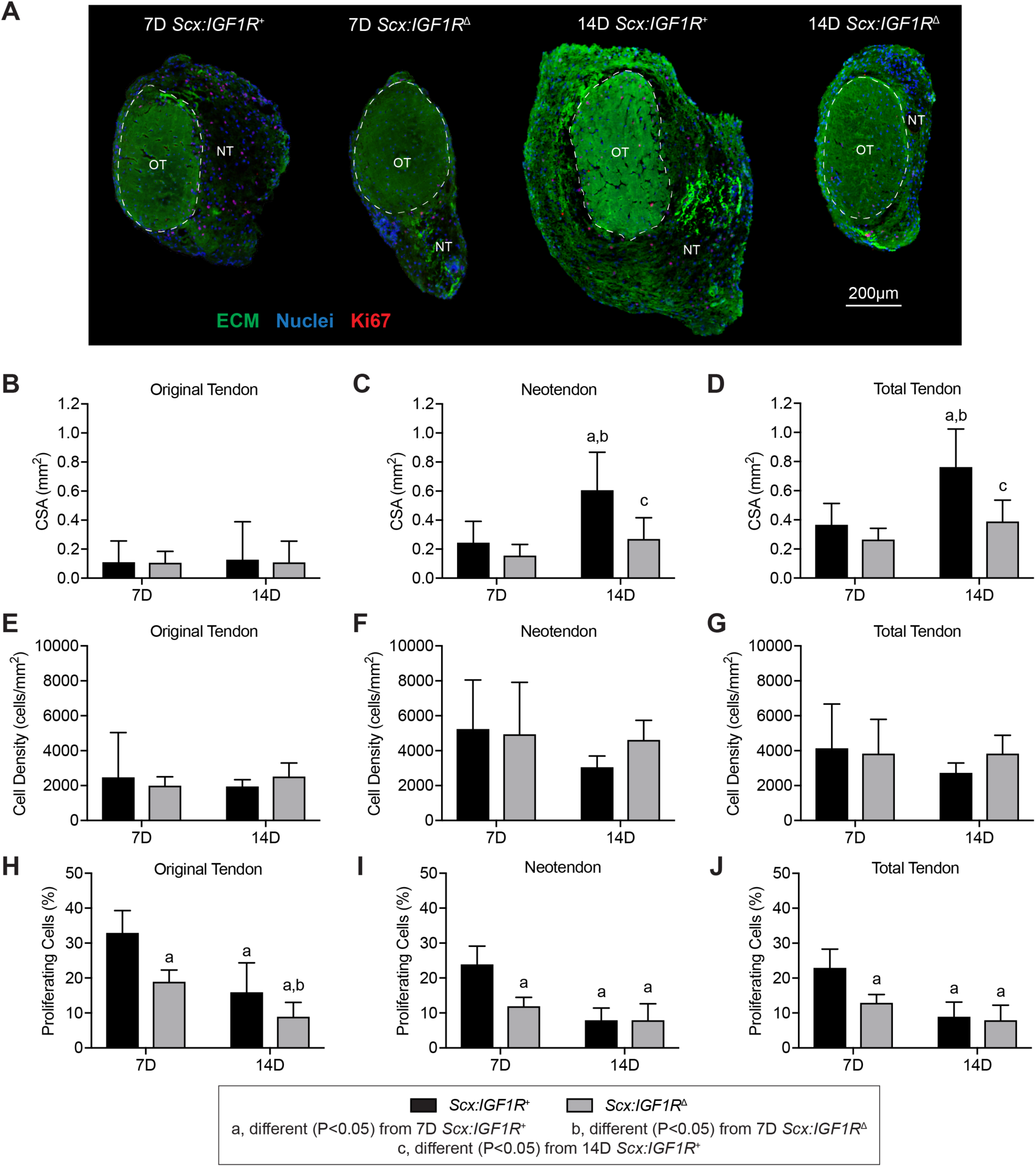
Effect of IGF1R deletion on tendon growth and cell proliferation. (A) Representative histology of cross-sections from *Scx:IGF1R*^+^ and *Scx:IGF1R*^Δ^ mice obtained at either 7 or 14 days after mechanical overload demonstrating general morphology, cell density, and abundance of proliferating cells. The original tendon (OT) and neotendon (NT) are indicated by the hashed line. Extracellular matrix (ECM), green; nuclei, blue; Ki67 (proliferating cells), red. Scale bar for all sections is 200µm. (B-D) Area measurements of tendons, with respect to the (B) original tendon, (C) neotendon, and (D) total tendon. (E-G) Cell density measurements of tendons, with respect to the (E) original tendon, (F) neotendon, and (G) total tendon. (H-J) Cell proliferation measurements, with respect to the (H) original tendon, (I) neotendon, and (J) total tendon. Values are mean±SD. Differences tested with a two-way ANOVA: a, significantly different (P<0.05) from 7D *Scx:IGF1R*^+^ mice; b, significantly different (P<0.05) from 7D *Scx:IGF1R*^Δ^ mice; c, significantly different (P<0.05) from 14D *Scx:IGF1R*^+^ mice. N≥4 mice per group.

We then sought to identify changes in the transcriptome of plantaris tendons of *Scx:IGF1R*^+^ mice and *Scx:IGF1R*^Δ^ mice in response to mechanical overload using RNAseq. At 7 days, there were 1108 transcripts that were at least 50% differentially regulated between genotypes, but only 159 transcripts at 14 days (Figure 3A-B). Pathway enrichment analysis was performed to evaluate cellular functions and signaling pathways predicted to be different between groups over time. Several of the pathways identified were involved with growth and differentiation, cytoskeletal signaling, and ECM production (Table 1). We selected several genes related to these processes to further explore and report in Figure 3C-E, and also performed qPCR validation of a subset of relevant genes (Table 2). Compared to non-overloaded controls, several growth factors and signaling molecules including *Adam12, Bmp1, Ctsd, Igf1, and Pappa* were upregulated in all overloaded groups, while *Bmp6, Fgf2, Inhbb*, and *Vegfa* were downregulated at all time points (Figure 3C). There was also an upregulation in *Bmp1*, *Bmp6*, *Igf1, Inhba*, *Pdgfa*, *Pdgfb*, *Tgfb1*, *Tgfb2*, *Wnt5b*, and *Wnt9a* in 7 day *Scx:IGF1R*^Δ^ mice compared to 7 day *Scx:IGF1R*^+^ mice, and by 14 days *Fgf2* and *Tgfb2* were significantly higher in *Scx:IGF1R*^Δ^ mice (Figure 3C). For genes involved in the liberation of IGF1 from IGFBPs, *C1s2* was upregulated at 7 days and *Ctsd* was upregulated at 14 days in *Scx:IGF1R*^Δ^ mice, while no differences in *Pappa* were observed between genotypes (Figure 3C). With regards to cell proliferation, and tenocyte specification and differentiation, *Acta2, Ccna2, Ccnb1, Ccne1, Cdk1, Cdk2, Cdk4, Cdk6, Cfi1, Itgav, Mki67, Pcna, Ptk2, Snai1*, *Trp53*, and *Vim* were upregulated in all overloaded groups (Figure 3D). *Cdkn1b, Itgb3, Rerg, S100a4, Scx, Snai1*, and *Yap1* were upregulated in 7 day *Scx:IGF1R*^Δ^ mice compared to 7 day *Scx:IGF1R*^+^ mice, while *Acta2, Cfl1, Itgav, Mcm6*, and *Pcna* were downregulated (Figure 3D). At 14 days, *Rerg* and *Scx* were upregulated in *Scx:IGF1R*^Δ^ mice (Figure 3D). For ECM genes, *Bgn, Co1a1, Co1a2, Col3a1, Col5a1, Col6a1, Col14a1, Fn1, Mmp2, Mmp3, Mmp14, Postn, Smoc2, Timp1, Tnc, Vcan*, and *Wisp1* were upregulated in overloaded groups, while *Comp* was downregulated (Figure 3E). In *Scx:IGF1R*^Δ^ mice, *Bgn, Col1a1, Col1a2, Col3a1, Col5a1, Comp, Fn1, Mmp2*, and *Mmp14* were upregulated at 7 days and *Col5a1* at 14 days, while *Mgp* was downregulated at 7 and 14 days compared to *Scx:IGF1R*^+^ mice (Figure 3E).

**Table 1.**
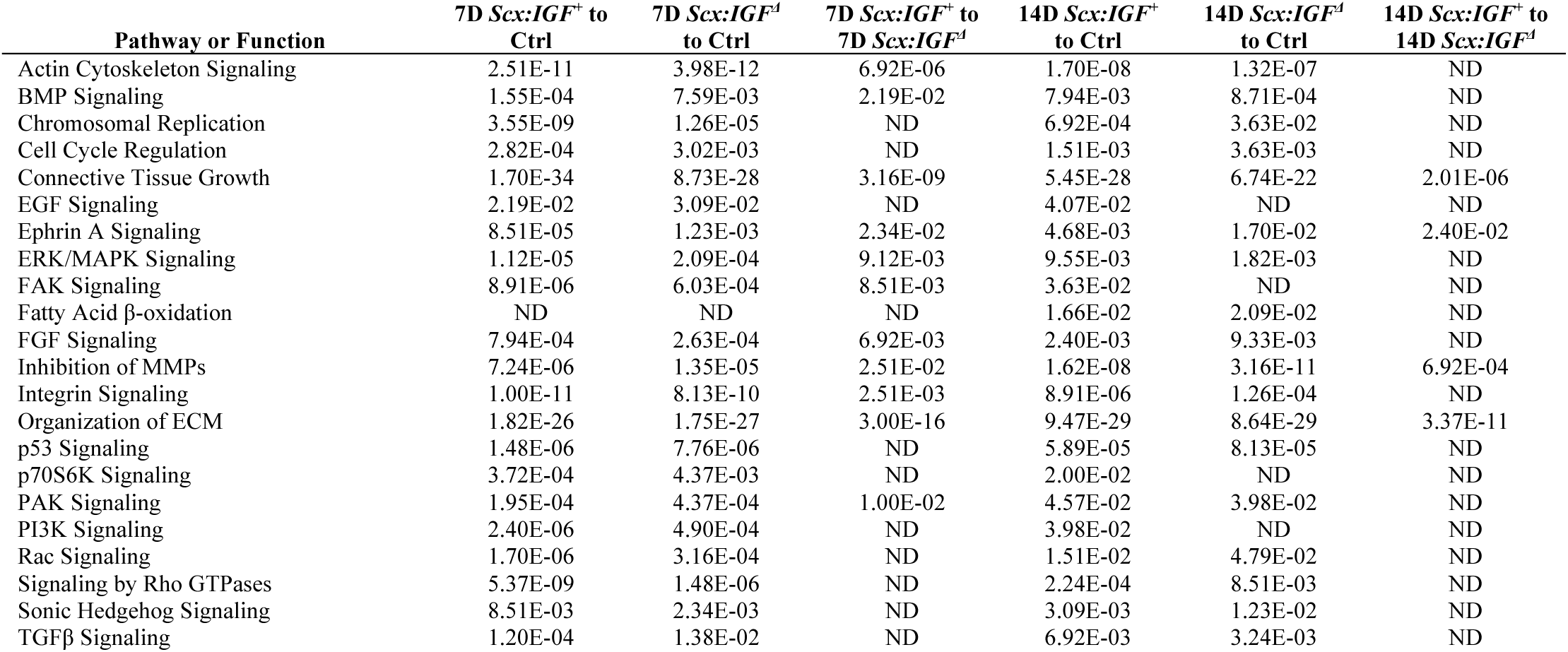
Gene enrichment analysis. q-values of select affected or differentially regulated pathways or biological functions that were identified using Ingenuity Pathway Analysis. ND, not significantly different.

**Table 2.** Gene expression data. Expression of genes as measured by qPCR. Values are mean±SD. Differences tested with a two-way ANOVA (α=0.05): a, different (P<0.05) from 7D *Scx:IGF1*^*+*^; b, (different (P<0.05) from 7D *Scx:IGF1*^*Δ*^; c, different (P<0.05) from 14D *Scx:IGF1*^*+*^.

| Gene | 7D *Scx:IGF*<sup>+</sup> | 7D *Scx:IGF1*<sup>Δ</sup> | 14D *Scx:IGF1*<sup>+</sup> | 14D *Scx:IGF1*<sup>Δ</sup> |
| --- | --- | --- | --- | --- |
| *CCNA2* | 8.19±1.90 | 4.48±0.66<sup>a</sup> | 3.88±1.27<sup>a</sup> | 4.38±0.97<sup>a</sup> |
| *CCNB1* | 3.09±0.43 | 1.81±0.18<sup>a</sup> | 1.19±0.60<sup>a</sup> | 1.43±0.36<sup>a</sup> |
| *CCND1* | 3.82±1.40 | 3.40±0.64 | 3.00±0.92 | 3.36±0.53 |
| *CCNE1* | 0.29±0.04 | 0.23±0.15 | 0.19±0.02 | 0.11±0.03<sup>a</sup> |
| *CDKN1A* | 2.07±0.39 | 3.04±0.85<sup>a</sup> | 2.23±0.46<sup>b</sup> | 1.74±0.34<sup>b</sup> |
| *COL1A1* | 224±72.4 | 372±46.7<sup>a</sup> | 482±133<sup>a</sup> | 561±74.5<sup>a,b</sup> |
| *COL1A2* | 889±316 | 1450±159<sup>a</sup> | 1890±551<sup>a</sup> | 2270±61.4<sup>a,b</sup> |
| *COL3A1* | 1308±346 | 1490±324 | 2070±640<sup>a</sup> | 2250±472<sup>a,b</sup> |
| *COL4A1* | 28.8±3.93 | 35.2±7.87 | 42.4±11.1<sup>a</sup> | 46.5±8.06<sup>a,b</sup> |
| *COL5A1* | 4.66±1.47 | 6.56±0.84 | 7.60±1.13<sup>a,c</sup> | 9.47±1.42<sup>a,b</sup> |
| *COL6A1* | 14.2±3.53 | 32.0±4.57<sup>a</sup> | 34.1±11.8<sup>a</sup> | 35.3±5.81<sup>a</sup> |
| *EGR1* | 2.36±0.97 | 1.75±0.88 | 1.16±0.50<sup>a</sup> | 0.93±0.45<sup>a</sup> |
| *EGR2* | 0.17±0.04 | 0.15±0.07 | 0.10±0.05<sup>a</sup> | 0.05±0.01<sup>a,b</sup> |
| *HAS2* | 2.19±0.34 | 0.79±0.11<sup>a</sup> | 1.20±0.42<sup>a</sup> | 0.93±0.23<sup>a</sup> |
| *MKI67* | 1.19±0.15 | 1.62±0.26 | 0.82±0.36<sup>a,b</sup> | 1.03±0.36<sup>a,b</sup> |
| *MKX* | 0.47±0.12 | 0.52±0.13 | 0.68±0.09<sup>a</sup> | 0.78±0.14<sup>a,b</sup> |
| *MMP2* | 5.37±1.11 | 7.95±1.53<sup>a</sup> | 11.2±2.17<sup>a,b</sup> | 11.4±1.57<sup>a,b</sup> |
| *MMP14* | 3.24±0.63 | 5.12±0.44<sup>a</sup> | 5.36±0.89<sup>a</sup> | 4.51±0.69<sup>a</sup> |
| *PCNA* | 22.2±4.63 | 14.3±3.84<sup>a</sup> | 11.9±1.48<sup>a</sup> | 13.8±2.61<sup>a</sup> |
| *SCX* | 9.97±2.25 | 98.9±15.6<sup>a</sup> | 19.3±7.62<sup>c</sup> | 265±107<sup>a,b</sup> |
| *TNMD* | 30.0±12.9 | 36.0±3.98 | 191±80.0<sup>a,b</sup> | 150±37.5<sup>a,b</sup> |

**Figure 3.**
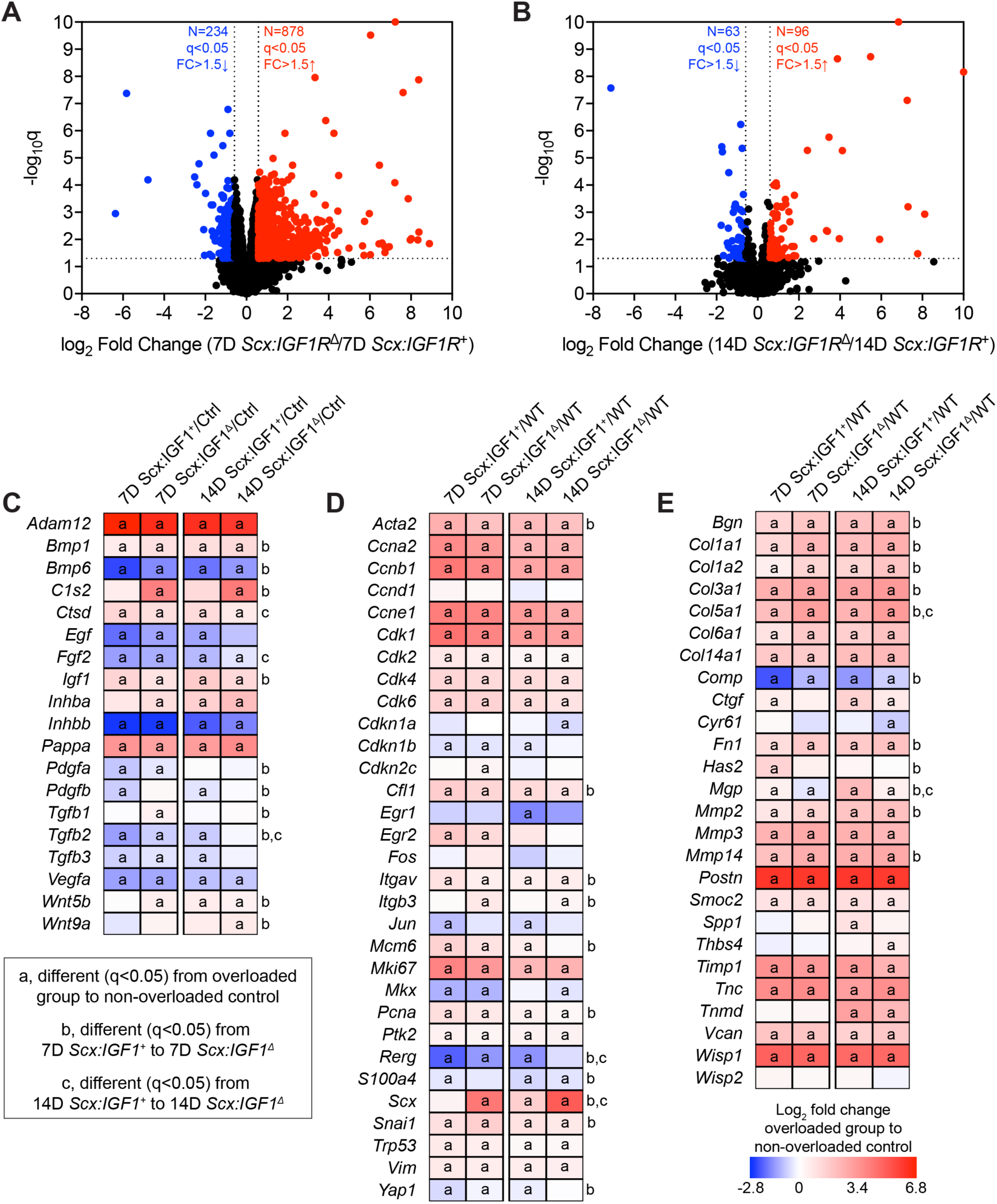
Whole tendon RNAseq. Volcano plot demonstrating log_2_ fold change (FC) and q values of all measured transcripts in plantaris tendons from *Scx:IGF1R*^+^ and *Scx:IGF1R*^Δ^ mice at (A) 7 days and (B) 14 days after surgical overload. Values greater than 10 are shown directly on the top or side border of the graph. Genes with a > 1.5-fold upregulation in *Scx:IGF1R*^Δ^ mice (log_2_ fold change > 0.585) and q value < 0.05 (-log10q > 1.301) are shown in red. Genes with a > 1.5-fold downregulation in *Scx:IGF1R*^Δ^ mice (log_2_ fold change < – 0.585) and q value < 0.05 (-log10q > 1.301) are shown in blue. All other genes are shown in black. (C-E) Heatmaps demonstrating the log_2_ fold change in selected genes from RNAseq that are (C) growth factors, cytokines, or genes involved with activating extracellular ligands, (D) involved in cell proliferation, and tenocyte specification and differentiation, or (E) components or regulators of the extracellular matrix. The fold change value is displayed for each group relative to non-overloaded *Scx:IGF1R*^+^ (Ctrl) mice. Differences between groups tested using edgeR: a, different (q<0.05) in the respective overloaded group to non-overloaded *Scx:IGF1R*^+^ mice; b, different (q<0.05) between 7D *Scx:IGF1R*^+^ and 7D *Scx:IGF1R*^Δ^ mice; c, different (q<0.05) between 14D *Scx:IGF1R*^+^ and 14D *Scx:IGF1R*^Δ^ mice. N=4 mice per group.

Based on the *in vivo* results which suggested a role for IGF1 in controlling cell cycle behavior and ECM synthesis involving PI3K/Akt and ERK signaling, we took a reductionist approach to evaluate IGF1 signaling *in vitro* using cultured primary tenocytes. We observed that IGF1 treatment resulted in IGF1R^Y1135^ phosphorylation, leading to early downstream activation of IRS1^Y608^, ERK1/2^T202/Y204^, Akt^T308^, Akt^S473^, p70S6K^T389^, and p70S6K^T421/S424^ (Figure 4A). While ERK1/2^T202/Y204^, Akt^T308^, and Akt^S473^ phosphorylation decreased following 5 minutes of IGF1 treatment, p70S6K^T389^ and p70S6K^T421/S424^ activation was sustained from 15 through 60 minutes after treatment, and phosphorylation of ELK1^S383^ was detected at 30 and 60 minutes (Figure 4A). Phosphorylation of the inhibitory IRS1^S612^ site was detected at 30 and 60 minutes following IGF1 treatment (Figure 4A).

**Figure 4.**
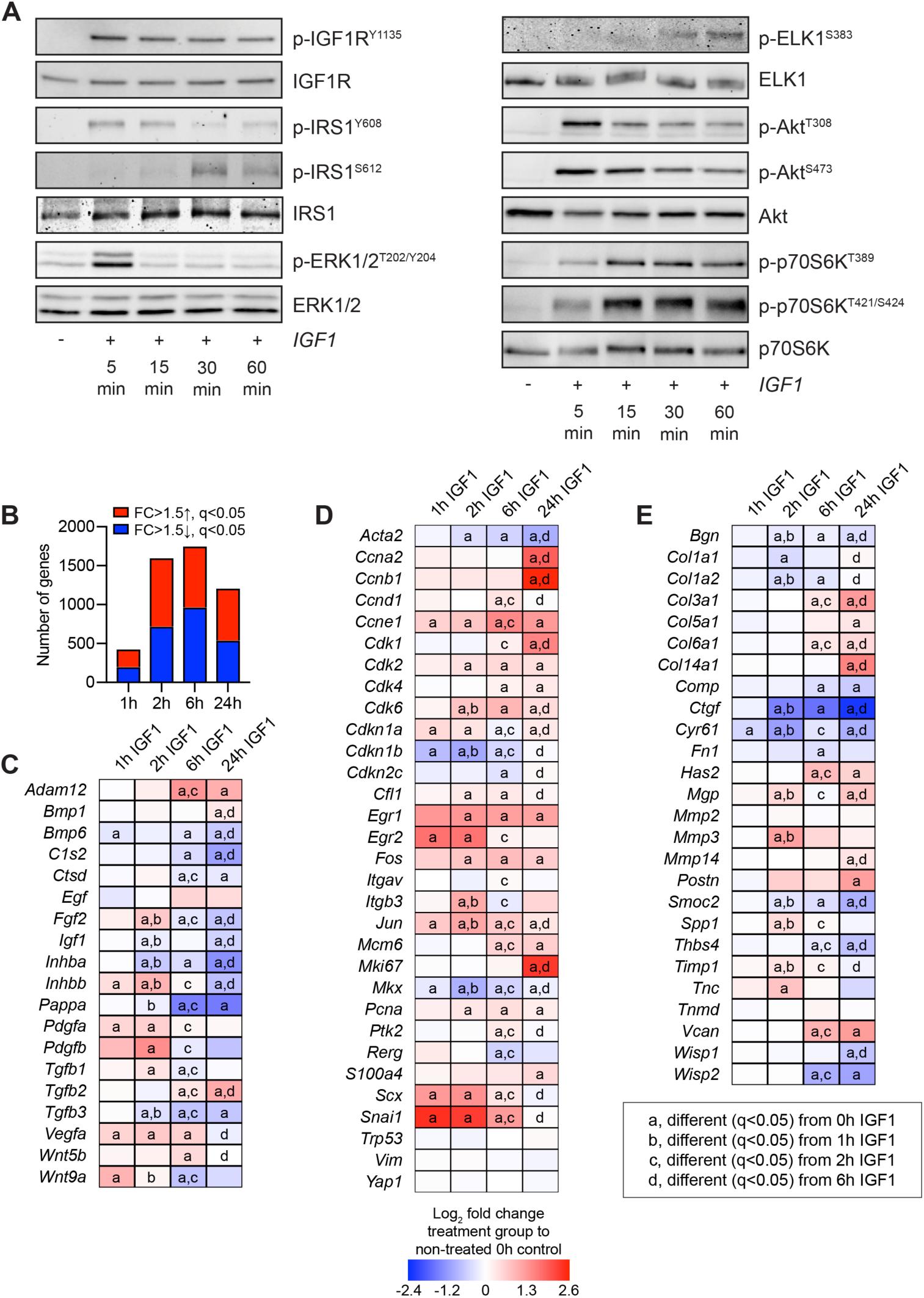
Signaling protein activation and RNAseq in cultured tenocytes treated with IGF1. (A) Representative western blots for p-IGF1R^Y1135^, total IGF1R, IRS1^Y608^, IRS1^S612^, total IRS1, p-ERK1/2^T202/Y204^, total ERK1/2, p-ELK^S383^, total ELK, p-Akt^T308^, p-Akt^S473^, total Akt, p-p70S6K^T389^, p-p70S6K^T421/S424^, and total p70S6K in cultured tenocytes that were untreated (0 min) or treated with IGF1 for 5, 15, 30, or 60 minutes. (B-E) RNAseq analysis of untreated tenocytes (0h), or tenocytes treated with IGF1 for 1, 2, 6, or 24 hours. The fold change (FC) value is displayed for each group is relative to tenocytes that were not treated with IGF1 (0 hours). (B) Number of genes that were significantly upregulated (red) with a greater than 1.5-fold upregulation and q < 0.05, and significantly downregulated (blue) with a greater than 1.5-fold upregulation and q < 0.05. (C-E) Heatmaps demonstrating the log_2_ fold change in selected genes from RNAseq that are (C) growth factors, cytokines, or genes involved with activating extracellular ligands, (D) involved in cell proliferation, and tenocyte specification and differentiation, or (E) components or regulators of the extracellular matrix. Differences between groups tested using edgeR: a, different (q<0.05) from 0h IGF1; b, different (q<0.05) from 1h IGF1; c, different (q<0.05) from 2h IGF1; c, different (q<0.05) from 6h IGF1.

To further explore the effect of IGF1 treatment on tenocytes, we performed RNAseq using cultured tenocytes that were not treated with IGF1, or treated with IGF1 for 1, 2, 6, or 24 hours. At 1 hour, over 400 transcripts were at least 50% differentially regulated and had a q<0.05, while over 1000 genes were similarly affected at 2, 6, and 24 hours (Figure 4B). We used the same panel of genes explored in whole tendons for further analysis in cultured tenocytes. Across these transcripts, *Egf, Mmp2, Tnmd, Trp53, Vim*, and *Yap1* were not affected by IGF1 treatment (Figure 4C-E). For the growth factors and signaling molecules that were differentially regulated in whole tendons of *Scx:IGF1R*^+^ and *Scx:IGF1R*^Δ^ mice, there was an induction of *Fgf2, Pdgfa, Pdgfb, Tgfb1*, and *Wnt9a* by 2 hours, while *Igf1* was downregulated compared to untreated cells (Figure 4C). By 24 hours, with the exception of *Bmp1* and *Tgfb2*, nearly all other growth factors and signaling molecules that were differentially regulated in tendons were downregulated in cultured tenocytes (Figure 4C). *Cfl1, Itgb3, Pcna, Scx*, and *Snai1* demonstrated an early upregulation in response to IGF1 treatment, while *Acta2* and *Cdkn1b* were initially reduced (Figure 4D). By 24 hours *Mcm6, Pcna*, and *S100a4* were upregulated and *Acta2* was suppressed (Figure 4D). The ECM genes *Mgp, Mmp3, Spp1, Timp1*, and *Tnc* were upregulated by 2 hours, while *Bgn, Col1a1, Col1a2, Ctgf, Cyr61*, and *Smoc2* were downregulated (Figure 4E). At 24 hours, *Col3a1, Col5a1, Has2, Mgp*, and *Mmp14* were induced, and *Bgn* and *Comp* were reduced in IGF1 treated tenocytes.

As we observed differences in cell proliferation *in vivo* and that IGF1 affected the expression of cell cycle control genes *in vitro*, we next determined if IGF1 directly impacts the proliferation of cultured tenocytes, and the role of the PI3K/Akt and ERK pathways in this process. We sought to validate the findings for *Mki67* expression in RNAseq data, and using qPCR we observed a slightly greater than 1.5-fold transient induction at 2 hours after IGF1 treatment, and a near 8-fold induction in *Mki67* at 24 hours (Figure 5A). In support of these findings, we observed a 2.5-fold increase in the number of BrdU^+^ tenocytes in response to IGF1 treatment (Figure 5B). Inhibiting the PI3K/Akt pathway with wortmannin did not impact IGF1-mediated tenocyte proliferation or *Mki67* expression, while blocking ERK1/2^T202/Y204^ with PD98059 reduced cell proliferation and *Mki67* expression (Figure 5B-C).

**Figure 5.**
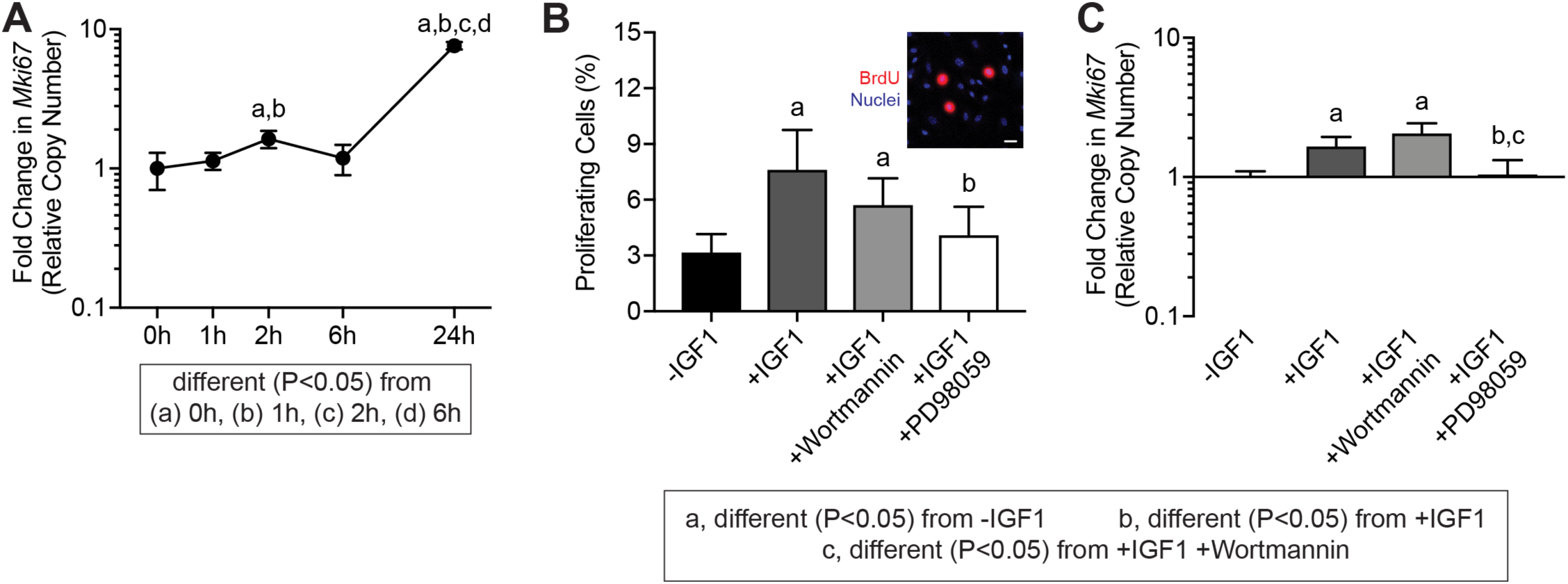
Proliferation in tenocytes treated with IGF1. (A) *Mki67* expression in untreated tenocytes (0h) or in tenocytes treated with IGF1 for 1, 2, 6, or 24 hours, measured with qPCR. Differences between groups tested using a one-way ANOVA: a, different (P<0.05) from 0h; b, different (P<0.05) from 1h; c, different (P<0.05) from 2h; d, different (P<0.05) from 6h. (B) The abundance of proliferating tenocytes (expressed as a percentage of total tenocytes) and (C) *Mki67* expression, in untreated cells or in cells treated with IGF1, IGF1 and wortmannin, or IGF1 and PD98059 for 2 hours. Inset (B) is a representative image demonstrating BrdU^+^ nuclei (red) and total nuclei (blue). Scale bar is 30µm. Differences tested with a one-way ANOVA: a, significantly different (P<0.05) from control; b, significantly different (P<0.05) from IGF1; c, significantly different (P<0.05) from IGF1 and wortmannin. Values are mean±CV (A,C) or mean±SD (B). N≥4 replicates per group.

Based on the differences in tendon size between *Scx:IGF1R*^+^ and *Scx:IGF1R*^Δ^ mice, the differential expression of *Col1a1* and *Col1a2* in whole tendons, and the cell culture RNAseq data, we sought to determine whether IGF1 directly induces type I collagen expression in tenocytes. Using qPCR, there were no differences in *Col1a1* and *Col2a2* expression following treatment with IGF1 (Figure 6A-B). We also did not observe dose- or time-dependent effects of IGF1 on type I collagen protein abundance in tenocytes, although we did note that cells that are in the S phase, as indicated by the presence of EdU^+^ nuclei, reduce type I collagen production (Figure 6C-E).

**Figure 6.**
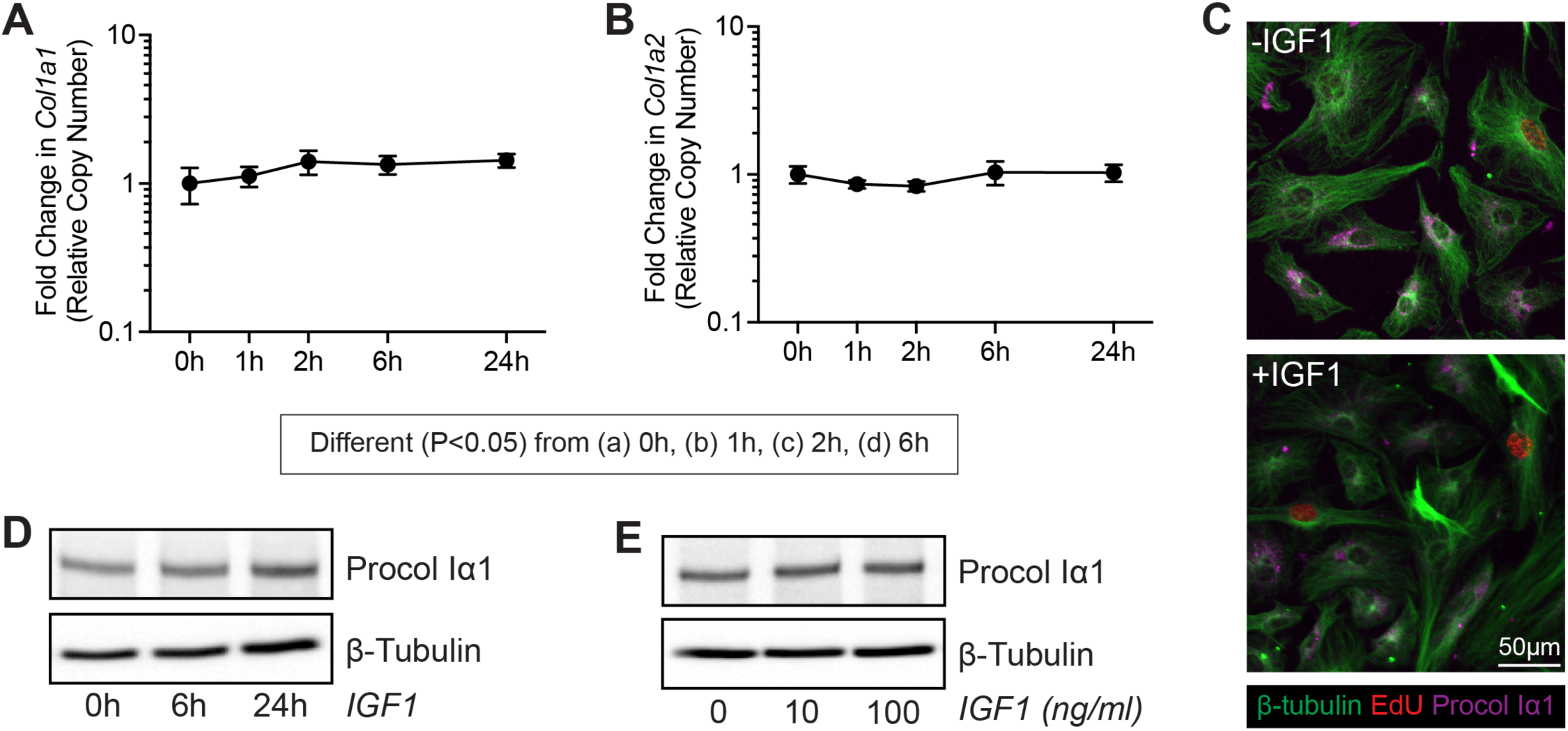
Type I collagen production in tenocytes treated with IGF1. (A) *Col1a1* and (B) *Col1a2* expression in untreated tenocytes (0h) or in tenocytes treated with IGF1 for 1, 2, 6, or 24 hours, measured with qPCR. Values are mean±CV. Differences between groups tested using a one-way ANOVA: a, different (P<0.05) from 0h; b, different (P<0.05) from 1h; c, different (P<0.05) from 2h; d, different (P<0.05) from 6h. (C) Representative immunocytochemistry of cultured tenocytes treated with 0ng/mL or 100ng/mL of IGF1 for 24 hours. β-tubulin, green; EdU, red; Procol Iα1, magenta. Scale bar is 50µm in all images. (D) Representative western blot for procollagen type Iα1 (Procol Iα1) in untreated tenocytes (0h) or tenocytes treated with 100ng/mL of IGF1 for 6 or 24 hours. (E) Representative western blot for Procol Iα1 in tenocytes treated with 0ng/mL, 10ng/mL or 100ng/mL of IGF1 for 24 hours.

Finally, we sought to determine the effect of IGF1 signaling on general protein synthesis. Using the SUnSET method in which the tyrosyl-tRNA analog puromycin is incorporated into newly synthesized proteins, we treated tenocytes with IGF1 and observed a 2-fold increase in puromycin incorporation. The broad spectrum translation inhibitor cycloheximide as well as wortmannin blocked IGF1-mediated protein synthesis in tenocytes, but surprisingly ERK1/2 inhibition resulted in a nearly 6-fold increase in protein synthesis (Figure 7A-B). To investigate this phenomenon in more detail, we probed for the presence of permissive and inhibitory phosphorylation sites on components of the PI3K/Akt and ERK1/2 signaling pathways. As anticipated PD98059 blocked ERK1/2^T202/Y204^ phosphorylation as well as phosphorylation of the downstream transcription factor ELK1^S383^, and wortmannin inhibited Akt^T308^ phosphorylation (Figure 7C). However, inhibition of ERK1/2 surprisingly increased Akt^T308^ phosphorylation independent of IRS1^Y608^ phosphorylation, suggesting ERK1/2 is acting to inhibit protein synthesis downstream of IRS1 (Figure 7C). Further, wortmannin decreased ERK1/2^T202/Y204^ and ELK1^S383^ phosphorylation and abolished phosphorylation of the inhibitory IRS1^S612^ site, leading to increased phosphorylation of the permissive IRS1^Y608^ site (Figure 7C). Based on these findings, we propose a model in which IGF1 regulates tenocyte proliferation and protein synthesis through coordinated and intersecting actions of the PI3K/Akt and ERK1/2 pathways (Figure 8).

**Figure 7.**
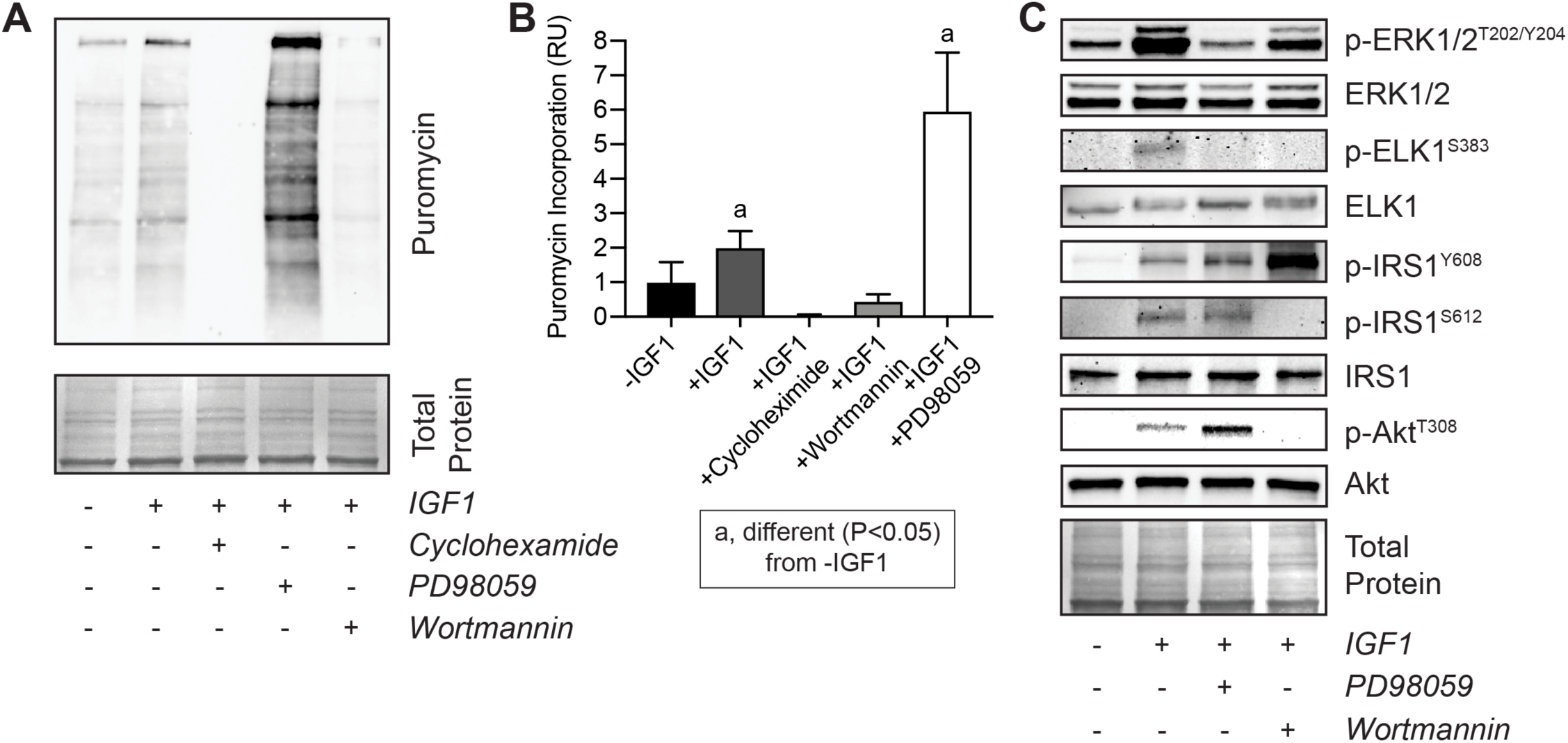
Protein synthesis in tenocytes treated with IGF1. (A) Representative western blot for proteins that have incorporated puromycin and (B) quantification of band densitometry for tenocytes that were untreated or were treated with IGF1, IGF1 and cycloheximide, IGF1 and wortmannin, or IGF1 and PD98059. (C) Representative western blots for p-ERK1/2^T202/Y204^, total ERK1/2, p-ELK^S383^, total ELK, IRS1^Y608^, IRS1^S612^, total IRS1, p-Akt^T308^, and total Akt in untreated cells or in cells treated with IGF1, IGF1 and wortmannin, or IGF1 and PD98059. Differences tested with a one-way ANOVA: a, significantly different (P<0.05) from -IGF1 controls. Coomassie stained membranes are shown as total protein loading controls. Values are mean±CV. N≥4 replicates per group.

**Figure 8.**
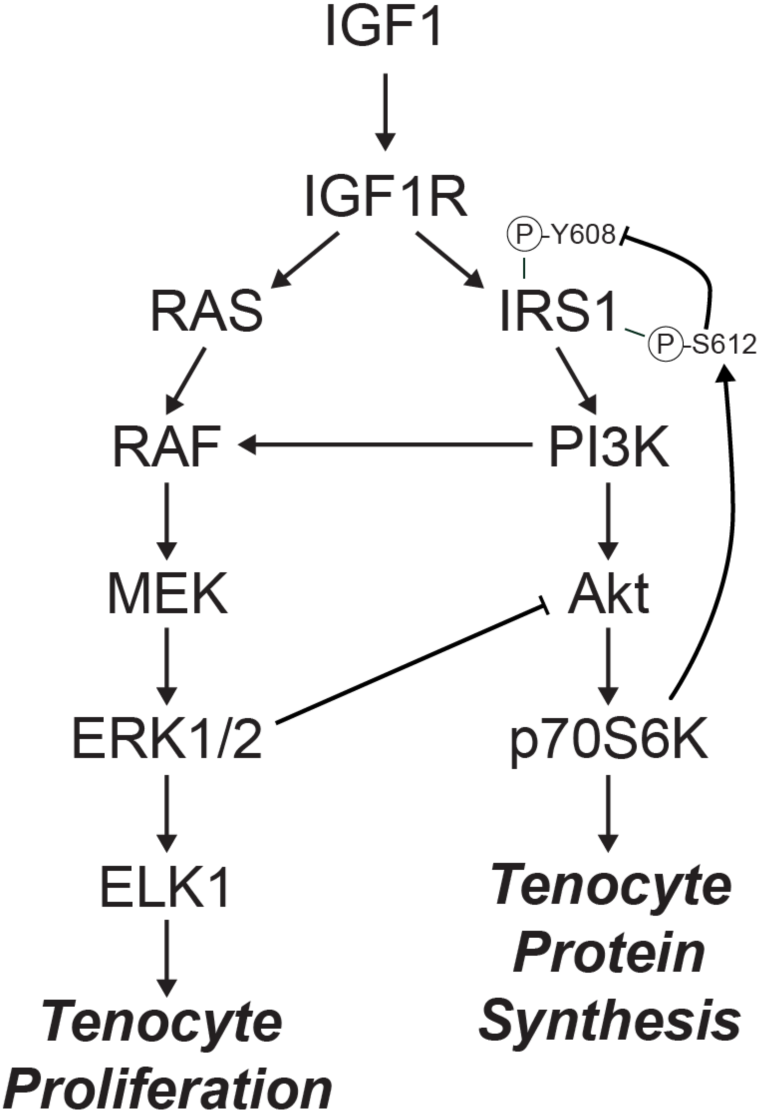
IGF1 signaling in tenocytes. Diagram of the proposed regulation of tenocyte proliferation and protein synthesis by IGF1.

## Discussion

IGF1 is known to play a critical role in the growth and adaptation of various musculoskeletal tissues, including skeletal muscle, bone, and cartilage, but less is known about tendon. To study tendon growth we used the synergist ablation technique, in which the Achilles tendon is surgically removed resulting in a supraphysiological growth stimulus to the synergist plantaris tendon and muscle (Gumucio et al., 2014; Hamilton et al., 2014; Mccarthy et al., 2011; Olesen et al., 2006; Schwartz et al., 2015; Sugg et al., 2017). A neotendon matrix consisting of immature collagen and other ECM proteins forms around the plantaris tendon, and over a one month period this ECM matures and resembles the original tendon matrix (Schwartz et al., 2015). By applying the synergist ablation model in a line of mice in which *IGF1R* was deleted in tenocytes, we demonstrated that IGF1 signaling is required for the proper growth of tendons in adult animals. Compared to *Scx:IGF1R*^+^ mice, the plantaris tendons of *Scx:IGF1R*^Δ^ mice had a reduced CSA and numbers of proliferating cells, and displayed alterations in growth factors, signaling molecules, cell cycle control genes, and ECM components. Using *in vitro* studies, we observed that IGF1 promotes a delayed increase in cell proliferation and directs protein synthesis, but surprisingly did not directly regulate type I collagen expression. IGF1 also induced the expression of *Egr1, Egr2*, and *Scx*, which are transcription factors that direct the expression of genes that are important for tendon development and growth (Léjard et al., 2011; Murchison et al., 2007), as well as *Snai1* which is involved in fibroblast-mediated tissue growth (Kalluri and Weinberg, 2009). Futher, we identified that both the PI3K/Akt and ERK pathways are activated downstream of IGF1, and work in a coordinated manner to regulate cell proliferation and protein synthesis. These results have important implications for our understanding of the basic biology of tendon growth and may inform the treatment of tendinopathies.

Numerous growth factors have been studied in the context of tenocyte proliferation *in vitro*. In the current study, we observed a minor increase in markers of cell proliferation within two hours of IGF1 treatment, but did not see notable increases until 24 hours. In mechanically stimulated tendons cell proliferation was reduced at 7 days in *Scx:IGF1R*^Δ^ mice. The CSA of tendons was similar between mice at 7 days, but there was no appreciable growth in the tendons of *Scx:IGF1R*^Δ^ mice between 7 and 14 days, while the tendons of *Scx:IGF1R*^+^ mice continued to grow thereafter. This suggests that IGF1 is regulating tenocyte proliferation in an indirect manner, perhaps through controlling the expression of other growth factors and signaling molecules that directly regulate cell cycle machinery. TGFβ1 signals through the TGFβRII and TGFβRI receptors to activate the Smad2/3 and MAPK signaling pathways in tenocytes (Gumucio et al., 2015), and treating tenocytes with recombinant TGFβ1 increases tenocyte proliferation (Mendias et al., 2012; Spindler et al., 1996). PDGFa and PDGFb, which are activated by mechanical loading and signal through the PDGFRα and PDGFRβ receptors that are members of the RTK family, also promote tenocyte proliferation and are required for proper tendon growth (Sugg et al., 2018; Thomopoulos et al., 2009). Related to PDGFa and PDGFb, FGF2 induces fibroblast growth and proliferation also through RTK signaling, and in human renal fibroblasts, treatment with TGFβ1 increased cell proliferation through induction of FGF2 expression (Strutz et al., 2001). In our data, the expression of *TGFβ1, PDGFa, PDGFb*, and *FGF2* was upregulated by 2 hours following IGF1 treatment in cultured tenocytes, which corresponded to modest increases in *Fos, Jun, Mki67*, *Pcna*, cyclin E (*Ccne1*), *Cdk2*, and *Cdk6*, along with a decrease in p27 (*Cdkn1b*). By 24 hours, *Bmp6* which inhibits cell proliferation (Arndt et al., 2019; Kersten et al., 2005) was downregulated, along with an upregulation in *Fos, Jun, Mki67* and *Pcna*. Several cyclins (*Ccna2, Ccnb1*, and *Ccne1*) and CDKs (*Cdk1, Cdk2, Cdk4*, and *Cdk6*) were also upregulated, and a downregulation in p27 was observed. At the whole tissue level, *Bmp6* was elevated and *Pcna* was decreased at 7 days in *Scx:IGF1R*^Δ^ mice compared to *Scx:IGF1R*^+^ mice, and several other growth factors and signaling molecules with direct roles in regulating cell proliferation were also differentially regulated. Therefore, it is likely that IGF1 chiefly acts in an indirect manner to regulate tenocyte proliferation through the regulation of other growth factors that act loacally. The delayed effect of IGF1 on enhancing tenocyte proliferation may explain why the size of the neotendon was not different between *Scx:IGF1R*^+^ and *Scx:IGF1R*^Δ^ mice at 7 days, but at 14 days the neotendon of the *Scx:IGF1R*^+^ mice was two-fold larger than *Scx:IGF1R*^Δ^ mice.

In addition to reducing cell proliferation, the loss of *IGF1R* in tenocytes resulted in smaller tendons of mechanically loaded mice. Since type I collagen is the major constituent of the tendon ECM, and previous studies have demonstrated that direct injection of IGF1 into tendons increases collagen synthesis (Hansen et al., 2013) and patients with acromegaly that have elevated levels of IGF1 also demonstrate an upregulation of *Col1a1* in their tendons (Doessing et al., 2010), we sought to determine whether IGF1 could directly regulate collagen synthesis in tenocytes. Based on these studies, we anticipated that IGF1 would directly induce type I collagen expression and that *Scx:IGF1R*^Δ^ mice would have reduced expression in their tendons in response to mechanical overload. However, *Col1a1* and *Col1a2*, as well as *Col3a1* and *Col5a1*, were upregulated in *Scx:IGF1R*^Δ^ mice compared to *Scx:IGF1R*^+^ mice, and treatment of tenocytes with IGF1 did not induce *Col1a1* or *Col1a2* expression and resulted in no change in type I procollagen protein abundance. These results indicate that IGF1 does not directly induce type I collagen gene expression nor increase type I collagen translation. In support of our observations of a delayed growth defect in tendons of *Scx:IGF1R*^Δ^ mice, 3-dimensional tissue engineered tendon constructs treated with IGF1 demonstrated increased size and hydroxyproline content, but this increase was not observed until at least two weeks after beginning treatment with IGF1 (Herchenhan et al., 2015). Numerous other genes that encode minor but important proteins that constitute or modify the ECM such as *Ctgf*, *Fn, Mmp2, Mmp3, Mmp14, Postn, Smoc2, Spp1, Thsb4, Timp1, Tnc, Tnmd, Vcan*, and *Wisp1* were induced by mechanical loading of tendons, and *Ctgf, Cyr61, Has2, Postn, Spp1, Thbs4, Timp1, Tnc*, and *Vcan* appeared to be directly or indirectly regulated by IGF1 in cultured tenocytes.

To further explore the difference in size between tendons of *Scx:IGF1R*^+^ and *Scx:IGF1R*^Δ^ mice, we determined whether IGF1 regulated general protein synthesis. IGF1 is known to activate the PI3K/Akt pathway which increases protein synthesis in skeletal muscle (Gumucio et al., 2015), and local IGF1 can increase protein fractional synthesis rate in tendons of elderly men and patients with Ehlers-Danlos syndrome (Nielsen et al., 2014b; 2014a). Using the SUnSET technique (Goodman and Hornberger, 2013; Goodman et al., 2011), we observed that treating tenocytes with IGF1 doubled protein synthesis rates, and blocking Akt phosphorylation returned protein synthesis levels to baseline. Based on these findings and our observations of IGF1 and type I collagen synthesis, we propose that the larger tendons in the *Scx:IGF1R*^+^ mice occurred in part due to an increase in overall protein synthesis compared to *Scx:IGF1R*^Δ^ mice, rather than a direct increase in collagen production. There are also other signaling molecules regulated by IGF1 or mechano-sensing pathways in whole tissue that influence collagen production. Further, the regulation of *Col1a1* and *Col1a2* translation is a complex process. While translation requires that active ribosomes bind to *Col1a1* and *Col1a2* mRNAs, translation initiation also requires the interaction of regulatory cofactors with the 5’ stem loop and 3’ untranslated region of the transcripts, and the binding of these transcripts to nonmuscle myosin and association with vimentin filaments (Stefanovic, 2013). Combined, our results indicate that IGF1 signaling contributes to protein synthesis in tenocytes through an Akt-dependent mechanism, but IGF1 does not appear sufficient to induce type I collagen translation and likely works in a coordinated fashion with other signaling pathways to regulate this process.

ERK is a well-known signaling pathway that is activated in response to mechanical loading, often through the integrin αV/β3/FAK pathway (Tahimic et al., 2013). In whole tendons, integrin αV (*Itgav*) and β3 (*Itgab3*) were induced in response to synergist ablation, and RNAseq pathway enrichment analysis predicted activation of integrin, FAK, and ERK pathways, with a differential response between *Scx:IGF1R*^+^ and *Scx:IGF1R*^Δ^ mice. IGF1 is known to work in coordination with integrin αV/β3/FAK signal transduction, with ERK as a common and important downstream effector kinase for both pathways (Tahimic et al., 2016). While mechanical loading is known to increase both ERK and PI3K/Akt activation in tendons (Havis et al., 2016; Scott et al., 2007; 2005; Sugg et al., 2018), the role of IGF1 in modulating ERK and PI3K/Akt in tendons was not known. In this study, we observed that IGF1 activates both ERK and PI3K pathways in tenocytes, and that ERK activation is required for the IGF1-mediated increase in tenocyte proliferation. ELK1 is a transcription factor downstream of ERK that directs the expression of genes involved in cell proliferation, such as *Fos* and *Jun* that subsequently regulate expression of cyclins and CDKs that control cell cycle progression (Bahrami and Drabløs, 2016; Boros et al., 2009; Cook et al., 1999; Liao et al., 1997). ELK1^S383^ was phosphorylated in response to IGF1 treatment, and as expected this process was blocked by inhibiting ERK^T202/Y204^ phosphorylation. We also observed that ELK1^S383^ phosphorylation was inhibited by blocking PI3K. In other cell types, the PI3K pathway can amplify ERK^T202/Y204^ phosphorylation through activating the MAP3K RAF(Ebi et al., 2013), and based on our results, there is likely a similar mechanism occurring in tenocytes.

In addition to promoting cell proliferation, IGF1 also increased protein synthesis by approximately two-fold in tenocytes, and this process was dependent upon PI3K/Akt activation. However, inhibiting ERK^T202/Y204^ phosphorylation resulted in a nearly six-fold induction in protein synthesis, which was also a phenomenon that was not anticipated since ERK activation is often associated with mild increases in protein synthesis. This lead us to look at potential interactions between the PI3K/Akt and ERK pathways. IGF1R phosphorylates IRS1^Y608^, which is the principal site of interaction between IRS1 and the SH2 domain of PI3K. This causes the localization of PI3K to the plasma membrane where it can catalyze the conversion of PIP_2_ into PIP_3_ (Gumucio and Mendias, 2013). Akt then binds membrane-bound PIP_3_ through an N-terminal pleckstrin homology (PH) domain, and is activated by PDK1. The recruitment of Akt to the membrane and the phosphorylation of Akt^T308^ through PDK1 allows Akt to be released into cytosol to activate mTOR, p70S6K, and other downstream effectors (Gumucio and Mendias, 2013). IRS1 can also be phosphorylated at S612, which blocks phosphorylation of its own Y608 residue, and therefore IRS1^S612^ phosphorylation inhibits the ability of IRS1 to activate PI3K (De Fea and Roth, 1997). In some cell types, p-ERK^T202/Y204^ can phosphorylate IRS1^S612^ (Andreozzi et al., 2004; De Fea and Roth, 1997), however we did not observe this response in tenocytes. Instead, p-ERK^T202/Y204^ appears to modulate protein synthesis downstream of IRS1, by inhibiting phosphorylation of Akt^T308^ either directly or by targeting a process downstream of IRS1^Y608^. This inhibitory role of ERK on the PI3K pathway explains the pronounced increase in protein synthesis when ERK signaling is inhibited. While p-ERK^T202/Y204^ did not play a role in IRS1^S612^ or IRS1^Y608^ phosphorylation, inhibition of Akt activation abolished IRS1^S612^ phosphorylation and increased IRS1^Y608^ phosphorylation, suggesting that Akt or a downstream effector molecule acts to negatively regulate the PI3K pathway at IRS1. Previous studies have indicated that p70S6K^T389^ can phosphorylate several inhibitory serine residues on IRS1 (Copps and White, 2012), and we propose that p70S6K^T389^ is acting in a similar way to phosphorylate IRS1^S612^ and provide feedback inhibition on IGF1-mediated PI3K/Akt activation in tenocytes. Combined, these results indicate that the PI3K/Akt and ERK pathways interact downstream of IGF1 to control tenocyte proliferation and protein synthesis.

There are several limitations to this study. We only included male mice, and additional experiments evaluating the role of sex will likely provide further insight into the role of IGF1 in tendon adaptation. The plantaris overload synergist ablation technique used in this study is a supraphysiological growth stimulus that exceeds the normal growth signals tendons experience with exercise training. We only evaluated through two weeks after mechanical overload, and it is likely that IGF1 continues to have an influence on neotendon matrix growth and maturation beyond this point. Tendon mechanical properties were not assessed in this study, which limits interpretations about functional changes in mechanically overloaded tendons. We focused our analysis on the PI3K/Akt and ERK pathways based on the bioinformatics results, but it is likely IGF1 also regulates other downstream pathways in tenocytes. *IGF1R* was not completely abolished in tendons as cells other than tenocytes may also express this receptor. Despite these limitations, we feel that this study provided novel insight into the role of IGF1 signaling in regulating tendon growth.

The ability of tendon to grow and respond to mechanical and biochemical cues requires the coordination of multiple biological processes. Studies of 3D tissue engineered tendon constructs in culture and exercise training in human subjects have shown that tendons grow best in response to intermittent mechanical loading with adequate rest periods built in between loading regimes (Couppé et al., 2008; Geremia et al., 2018; Huisman et al., 2014; Paxton et al., 2012; Svensson et al., 2016). Failure of tendons to respond to these cues can lead to the development of tendinopathies or tendon ruptures (Mead et al., 2018). The results in the current study demonstrate an important role for IGF1 signaling in the growth of tendons in adult animals, and provide mechanistic support for the potential use of IGF1 in the treatment of tendinopathies. This is further supported by results from a small trial that demonstrated IGF1 increased collagen content and cell proliferation in horses with tendinopathy (Dahlgren et al., 2002), and local injection of IGF1 into the tendons of elderly men increased local collagen production (Nielsen et al., 2014b). However, it is unlikely that the use of growth factors alone would be sufficient to treat tendon disorders. High load eccentric resistance training can reduce pain and improve ECM structural organization in patients with tendinopathy, although this training does not lead to full symptomatic resolution for many patients (Kongsgaard et al., 2009; 2010). Recent studies have demonstrated that tendons synthesize collagen during rest phases at night, and assemble collagen into mature fibrils during the active day phases (Pickard et al., 2019; Yeung and Kadler, 2019). Combining the use of recombinant growth factors with properly timed mechanical loading interventions and appropriate rest periods could lead to the improved treatment of tendon disorders. While IGF1 appears to be a critical pathway in modulating tendon growth, further studies that integrate molecular genetics, cell biology, and mechanotransduction will provide additional insight into the basic mechanisms of tendon growth and likely lead to improved treatments of painful and debilitating tendon disorders.

## Methods

### Mice

All animal studies were approved by the University of Michigan and Hospital for Special Surgery/Weill Cornell Medical College/Memorial Sloan Kettering Institutional Animal Care & Use Committees. We used three strains of mice in this study. Wild type C57BL/6J (strain 000664) mice, and *IGF1R*^flox^ mice (strain 012251) in which exon 3 of *IGF1R* is flanked by loxP sites (Dietrich et al., 2000), were obtained from the Jackson Laboratory (Bar Harbor, ME, USA). *Scx*^CreERT2^ mice in which an *IRES-CreERT2* sequence was inserted between the stop codon and 3’ UTR in exon 2 of scleraxis (Howell et al., 2017), were kindly provided by Dr. Ronen Schweitzer (Shriners Hospitals for Children, Portland, OR, USA). Genotype of mice was determined by PCR analysis of DNA obtained from a tail tendon biopsy. After performing initial crosses between *Scx*^CreERT2/CreERT2^ and *IGF1R*^flox/flox^ mice, we generated *Scx*^CreERT2/CreERT2^:*IGF1R*^flox/flox^ mice to allow us to inactivate *IGF1R* in scleraxis expressing cells upon treatment with tamoxifen (referred to as *Scx:IGF1R*^Δ^ mice), while *Scx*^+/+^:*IGF1R*^flox/flox^ mice would maintain the expression of *IGF1R* in scleraxis expressing cells after tamoxifen treatment (referred to as *Scx:IGF1R*^+^ mice).

### Synergist Ablation Tendon Growth Procedure

Mice were treated daily with an intraperitoneal injection of 1mg of tamoxifen (T5648, Sigma, St. Louis, MO, USA) dissolved in 50 µl of corn oil to activate CreERT2 recombinase in scleraxis-expressing cells. Tamoxifen treatment began 3 days prior to surgery, and continued on a daily basis until tissue was harvested for analysis. Bilateral synergist ablation procedures were performed under isoflurane anesthesia as described previously (Gumucio et al., 2014; Sugg et al., 2018). An overview of the time points and surgical procedures are shown in Figure 1B-C. The Achilles tendon was surgically excised to prevent the gastrocnemius and soleus muscles from plantarflexing the talocrural joint, resulting in compensatory hypertrophy of the adjacent synergist plantaris muscle and tendon. A small incision was created in the skin above the posterior paw plantarflexor tendons, and the Achilles tendon was isolated and excised, leaving stumps at the myotendinous junction and calcaneus. The skin was closed with GLUture (Zoetis, Kalamazoo, MI, USA), buprenorphine was administered for post-operative analgesia, and *ad libitum* weight-bearing and cage activity were allowed in the postoperative period. Mice were closely monitored during the postoperative period for any adverse reactions. At tissue harvest, the left plantaris tendons were divided into proximal and distal halves, and snap frozen at −80ºC for gene expression analysis, while the right plantaris tendons were used for histology. After the tendons were removed, mice were euthanized by cervical dislocation. Plantaris tendons from additional non-overloaded *Scx:IGF1R*^+^ mice were obtained as described above for gene expression analysis.

### Histology

Histology was conducted as previously described (Gumucio et al., 2014; Sugg et al., 2018). Plantaris tendons obtained from animals were immediately placed in 30% sucrose solution for one hour, snap frozen in Tissue-Tek OCT Compound (Sakura Finetek, Torrance, CA, USA) and stored at −80°C until use. Tendons were sectioned at a thickness of 10 µm in a cryostat. Sections were stained with hematoxylin and eosin (H&E) to determine tendon cross-sectional area (CSA) and cell density. To evaluate proliferating cells, tendon sections were fixed in 4% paraformaldehyde, permeabilized with 0.2% Triton X-100, blocked with 5% goat serum, and incubated with rabbit anti-Ki67 antibodies (1:100; ab16667, Abcam, Cambridge, MA, USA) and goat anti-rabbit antibodies conjugated to AF555 (1:300; A-21429, Thermo Fisher Scientific), as well as wheat germ agglutinin (WGA) lectin conjugated to Alexa Fluor 488 (AF488) (1:200; W11261, Thermo Fisher Scientific, Carlsbad, CA, USA) to identify the extracellular matrix, and DAPI (1:500; Sigma) to label nuclei. Slides were imaged using an BX-51 microscope and camera (Olympus, Center Valley, PA, USA) for the H&E slides, while a A1 confocal laser microscope (Nikon Instruments, Tokyo, JP) was used for the Ki67 slides. Quantification of tendon size and cell density was performed using ImageJ (NIH, Bethesda, MD, USA).

### Cell Isolation

Tenocytes were isolated from the tail tendons of mice as described previously (Hudgens et al., 2016; Sugg et al., 2018). Mice were deeply anesthetized with isoflurane, the tail was removed, and animals were euthanized by cervical dislocation. Fascicles of tail tendons were isolated, finely minced and placed in low-glucose Dulbecco’s Modified Eagle Medium (DMEM; Gibco, Carlsbad, CA, USA) containing 0.2% type II collagenase (Gibco) for one hour at 37°C with constant agitation. An equal volume of growth medium (GM), which contains low-glucose DMEM with 10% fetal bovine serum (FBS; Gibco) and 1% antibiotic-antimycotic (AbAm; Gibco), was added to the digested tissue to inactivate the collagenase. Tenocytes were pelleted by centrifugation at 2500*g* for 10 minutes, resuspended in GM and plated. All dishes or chamber slides used in the study were coated with type I collagen (Corning, Corning, NY, USA). Cells were maintained in humidified incubators at 37°C and 5% CO_2_. Passage 2-4 tenocytes were used in experiments.

### In Vitro IGF1 Cell Culture Time Course

Tenocytes were grown to 60% confluence in GM, and switched to medium containing DMEM with 2% horse serum (Gibco) and 1% antibiotic-antimycotic (Gibco), referred to as differentiation medium (DM) overnight. The next day, media was replaced with DM containing 100ng/mL of IGF1 (R&D Systems, Minneapolis, MN, USA) for a period of time ranging from 1, 2, 6, or 24 hours. Cells that underwent the same procedure but did not receive IGF1 treatment are referred to as 0 hour cells. At the end of the treatment period RNA was isolated as described below.

### In Vitro Signal Transduction Assays

Tenocytes were grown to 90% confluence in GM and serum starved in DMEM with 1% AbAm for three hours, followed by treatment with either the mitogen-activated protein kinase kinases 1 and 2 (MEK1/2) inhibitor PD98059 (50 μM; InvivoGen, San Diego, CA, USA) to block ERK1/2 activation, or the PI3K inhibitor wortmannin (10 μM; InvivoGen) to inhibit Akt activation for 1 hour. Cells were then treated with 100ng/mL IGF1 for 5, 15, 30, or 60 minutes, scraped from their dishes and homogenized in RIPA buffer (Thermo Fisher Scientific) containing 1% protease and phosphatase inhibitors (Thermo Fisher Scientific).

### In Vitro Proliferation

Cell proliferation as measured by the uptake of bromodeoxyuridine (BrdU) was measured as previously described (Sugg et al., 2018). Tenocytes at 50% confluence were incubated overnight in DM, and then treated with DM containing 100 ng/ml of IGF1 (R&D Systems, Minneapolis, MN, USA), PD98059 (50 μM; InvivoGen), or wortmannin (10 μM; InvivoGen). Following a 16 hour overnight incubation, fresh media was added along with 20 μM of BrdU (Sigma) for one hour. After treatment with BrdU, cells were fixed in ice-cold methanol, permeabilized with 0.5% Triton X-100, and the BrdU epitope was exposed by denaturing DNA with 2 M HCl. Cells were then incubated with anti-BrdU antibodies (1:50; G3G4, Developmental Studies Hybridoma Bank, Iowa City, IA, USA) and secondary antibodies conjugated to AF555 (1:200; A-21127, Thermo Fisher Scientific), and DAPI (1:500) to identify nuclei. The number of BrdU^+^ nuclei as a fraction of total nuclei were quantified in five randomly selected fields of a single experiment from four independent experiments performed per group. Plates were imaged in an EVOS FL microscope (Thermo Fisher Scientific) and quantified with ImageJ software.

### In Vitro SUnSET Labeling

Protein synthesis in cultured tenocytes was measured using a surface sensing of translation (SUnSET) technique, as modified from studies in C2C12 myoblast cells (Goodman et al., 2011). Tenocytes were grown to 90% confluence in GM and serum starved in DMEM with 1% AbAm for three hours, followed by treatment with either PD98059, wortmannin, or the protein synthesis inhibitor cycloheximide (R&D Systems) for one hour. Cells were then treated for 30 minutes with 0.25µM puromycin (MilliporeSigma, Burlington, MA, USA), which is a tyrosyl-tRNA analog that is incorporated into newly translated proteins (Goodman and Hornberger, 2013), followed by 100ng/mL of IGF1 for one hour. At the end of the treatment period, tenocytes were scraped from their dishes and homogenized in RIPA buffer (Thermo Fisher Scientific) containing 1% protease and phosphatase inhibitors (Thermo Fisher Scientific).

### In Vitro Procollagen I Labeling

To label procollagen I in proliferating or non-proliferating cells, tenocytes were cultured as described above, incubated with DM overnight, and then treated with either normal DM or DM containing 100ng/mL of IGF1. The thymidine analog EdU was used in lieu of BrdU, as the acid denaturing step required for BrdU detection degraded procollagen I epitopes. Following a 24 hour incubation, fresh media was added along with 10 µM of EdU (Thermo Fisher Scientific) for one hour. Cells were fixed in 4% paraformaldehyde, permeabilized with 0.5% Triton X-100, and EdU was detected using a Click-iT kit (Thermo Fisher Scientific). Cells were then incubated with antibodies against procollagen I (1:100; sc-30136, Santa Cruz Biotechnology, Santa Cruz, CA, USA) and β-tubulin (1:200; ab6161, Abcam) and secondary antibodies conjugated to AF488 or AF647 (1:300; A11006 and A32733, Thermo Fisher Scientific). Slides were imaged in a LSM 880 laser scanning confocal microscope (Zeiss, Thornwood, NY, USA).

### RNA Sequencing and Gene Expression

RNA was isolated from tendons and cultured tenocytes as previously described (Schwartz et al., 2015; Sugg et al., 2018). Plantaris tendons or tenocytes were homogenized in QIAzol (Qiagen, Valencia, CA, USA) and RNA was purified with a miRNeasy Micro Kit (Qiagen) supplemented with DNase I (Qiagen). RNA concentration and quality was determined using a NanoDrop 2000 (Thermo Fisher Scientific) and a TapeStation 2200 (Agilent Technologies, Santa Clara, CA, USA).

For RNAseq, 250ng (for whole tendons) or 500ng (for culture cells) was delivered to the University of Michigan Sequencing Core for analysis. Sample concentrations were normalized and cDNA pools were created for each sample, and then subsequently tagged with a barcoded oligo adapter to allow for sample specific resolution. Sequencing was carried out using an Illumina HiSeq 2500 platform (Illumina, San Diego, CA, USA) with 50bp single end reads. Raw RNAseq data was quality checked using FastQC v0.10.0 (Barbraham Bioinformatics, Cambridge, UK). Alignment to the reference genome (mmu10, UCSC), differential expression based on counts per million mapped reads (CPM), and post-analysis diagnostics were carried out using edgeR (Robinson et al., 2010). A Benjamini-Hochberg false discovery rate (FDR) procedure was applied to correct P values for multiple observations, and these FDR-corrected P values are reported as q values. Sequencing data has been deposited to NIH GEO (GSE131804).

For qPCR, using iScript Reverse Transcription reagents (Bio-Rad, Hercules, CA, USA), RNA was reverse transcribed into cDNA which was amplified in a CFX96 real-time thermal cycler (Bio-Rad) using iTaq Universal SYBR Green Supermix (Bio-Rad). Target gene expression was normalized to the stable housekeeping gene cyclophilin D (*Ppid*) using the 2^−ΔCt^ method. Cyclophilin D was selected as a housekeeping gene from RNAseq data and validated with qPCR. For cell culture experiments, relative copy number was calculated using the linear regression of efficiency method (Rutledge and Stewart, 2010). Primer sequences are provided in Supplemental Material 1.

### Biological Pathway Analysis

Expression data from RNAseq measurements was imported into Ingenuity Pathway Analysis (IPA) software (Qiagen) to assist in predicting cellular and molecular pathways and processes that were altered in response to manipulating IGF1 signaling in tendons.

### Western blots

Western blots were performed as described previously (Gumucio et al., 2019; Sugg et al., 2018). Tendons and cell pellets were homogenized in ice cold RIPA buffer (Thermo Fisher Scientific) supplemented with 1% protease and phosphatase inhibitor cocktail (Thermo Fisher Scientific). A BCA assay (Thermo Fisher Scientific) was used to determine protein concentration. Protein homogenates were diluted in Laemmli’s sample buffer, placed in boiling water for two minutes, and 20 μg of protein was separated on either 6% or 12% SDS-PAGE gels depending on the protein of interest. Proteins were transferred to nitrocellulose or PVDF membranes (Bio-Rad) using the Trans-Blot SD semi-dry transfer apparatus (Bio-Rad), blocked with 5% non-fat powdered milk in TTBS solution and incubated with primary antibodies from Cell Signaling (1:1000; Danvers, MA, USA) against p-IGF1Rβ^Y1135^ (3918), IGF1Rβ (3025), p-IRS1^S612^ (2386), IRS1 (2382), p-Akt^T308^(13038), p-Akt^S473^ (4060), Akt (4691), p-p70S6K^T389^ (9234), p-p70S6K^T421/S424^ (9204), p70S6K (2708), p-ERK1/2^T202/Y204^ (4370), ERK1/2 (4695), or primary antibodies from Santa Cruz Biotechnology (1:1000) against procollagen type I (sc-30136), or primary antibodies from Abcam (1:1000) against β-tubulin (ab6046), p-ELK^S383^ (ab34270) or ELK (ab32106), or primary antibodies from MilliporeSigma against pIRS1^Y608^ (1:1000) or puromycin (1:2000; 12D10). Following primary antibody incubation, membranes were rinsed and incubated with HRP-conjugated secondary antibodies (1:10,000; from either AbCam or Cell Signaling). Proteins were detected using Clarity enhanced chemiluminescent reagents (Bio-Rad) and visualized using a digital chemiluminescent documentation system (Bio-Rad). Coomassie staining of membranes was performed to verify equal protein loading.

### Statistics

Results are presented as mean±standard deviation (SD) or mean±coefficient of variation (CV). Prism version 8.0 (GraphPad Software, La Jolla, CA, USA) was used to conduct statistical analyses for all data except RNAseq. A two-way ANOVA followed by Fisher’s *post hoc* sorting (α=0.05) evaluated the interaction between time after synergist ablation and *IGF1R* knockdown. For cell culture experiments, differences between groups were tested with an unpaired Student’s t-test (α=0.05) or a one-way ANOVA followed by Fisher’s *post hoc* sorting (α=0.05).

## Acknowledgements

This work was supported by NIH grants R01-AR063649 and F32-AR067086. The *Scx*^CreERT2^ mice in this report were kindly provided by Dr. Ronen Schweitzer of Shriners Childrens Hospital of Portland. We would like to acknowledge technical contributions from Dr. David Oliver at the Hospital for Special Surgery and Dr. James Markworth at the University of Michigan.

## Author Contributions

NPD, KBS, and CLM designed research; NPD, KBS, JRT, DCS, BJR, and CLM performed research; NPD, KBS, and CLM analyzed data; NPD, KBS, and CLM wrote the paper.

## Supplemental Material

**Supplemental Material S1.**
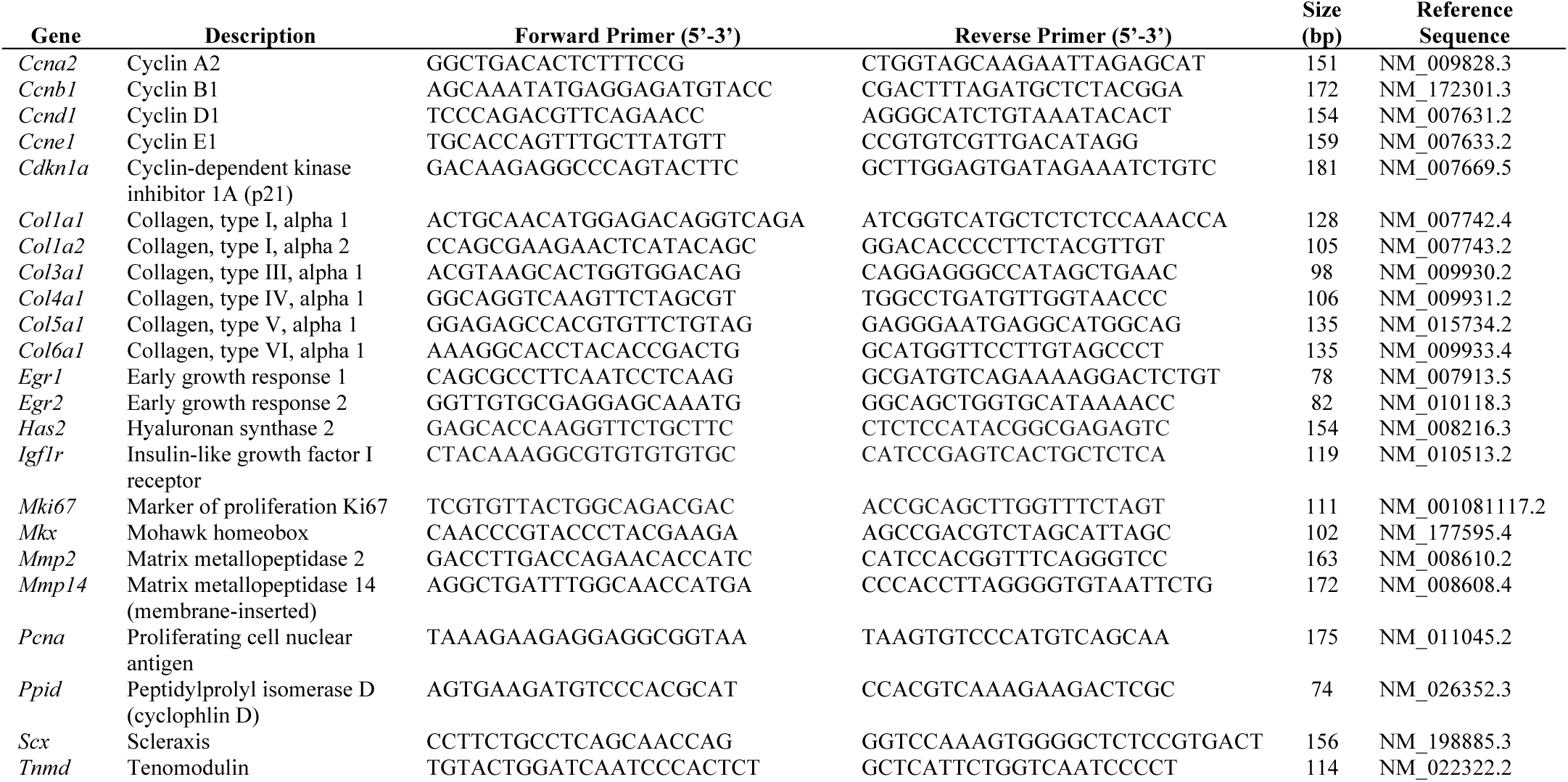
Primer sequences used for quantitative PCR.

